# AbSolution and ENCORE: a proof-of-concept for automating computational reproducibility in interactive applications

**DOI:** 10.64898/2026.05.16.724131

**Authors:** Rodrigo García-Valiente, Samuel H. Langton, Antoine H.C. van Kampen

## Abstract

Reproducibility and transparency in computational analyses are essential in science, although achieving these goals often requires significant knowledge and systematic organization. Graphical interactive applications simplify the conduct of analyses and make them accessible to a broader audience. However, there is currently no consensus on how to and to which extent implement reproducibility in interactive applications. We recently developed AbSolution, a user-friendly and flexible interactive web-based R Shiny application for exploring immune repertoires and their sequence-based features, and we established the ENCORE framework to enhance transparency and reproducibility by guiding researchers in structuring and documenting computational projects. In this work, as a proof-of-concept we integrate AbSolution, ENCORE and specific R packages to address reproducibility challenges. This enables a single-step export of raw, processed, and meta-data, the software environment, the underlying generated code and a HTML report containing results and figures, operating system, hardware and R session details, and researcher notes. Its reproducibility has been independently validated by the CODECHECK initiative. This paper demonstrates how the combination of several approaches can improve and automate reproducibility of interactive applications.

## Introduction

Lack of reproducibility of analyses and results remains a growing concern in scientific research (Dudda et al., 2025; Ioannidis, 2005; Stupple et al., 2019). Here, computational reproducibility refers to the ability to re-execute the same analysis using the original data, code, software environment, and parameters to obtain identical results. Ideally, the experimental and computational constituents of a research project are transparent and reproducible to improve the trustworthiness and reusability of research outcomes (Diaba-Nuhoho & Amponsah-Offeh, 2021; Turkyilmaz-van der Velden et al., 2020). However, this generally requires the research team to spend additional time and effort (Allen & Mehler, 2019). In addition, it may require further training to become acquainted with best practices (Brito et al., 2020). Specifically, reproducibility of computational analyses has been addressed by numerous research and initiatives. For example, guidelines to improve reproducibility (Brito et al., 2020; Sandve et al., 2013), the use of virtual computing environments for software versions, containers, and virtual machines (Gruning et al., 2018; Hunt & Gagnon-Bartsch, 2023), the use of git software versioning (Ram, 2013), and the importance of software engineering training (Connolly et al., 2023). Initiatives as the CODECHECK project (Nust & Eglen, 2021) and EVERSE (https://everse.software/) facilitate improving executability and reproducibility of computations.

One particular challenge involves the reproducibility of computational analyses performed using interactive applications. An interactive application is a software tool that enables to engage directly with data and data analysis approaches through reactive programming and a graphical (web-based) user interface. It allows real-time input and feedback, facilitating dynamic exploration, visualization, and analysis of results without the need for manual code execution. The well-known Galaxy platform addresses this issue by automatically tracking software versions, initial inputs, objects and parameters for its hosted applications and tools ecosystem (Galaxy, 2022). In addition, Galaxy further enhances reproducibility through its workflow engine, which supports the sharing and reuse of workflows and associated analysis objects. However, even within such platforms, interactive tools typically make reproducibility difficult by not recording the steps taken during analysis (Gadhave et al., 2022), particularly manual interactions, by hiding their domain logic -that is, the application-specific analytical rules and data transformations- and by introducing additional code complexity. For the widely-adopted Shiny applications (Chang et al., 2024), packages such as *shinymeta* (Cheng & Sievert, 2025) can help expose this domain logic by capturing it to generate the corresponding executable code.

Availability of data and code alone is not always sufficient to ensure reproducibility(Mendes, 2018; Papin et al., 2020; Stodden et al., 2018; Tiwari et al., 2021) but it may also require the use of containerized environments(Hunt & Gagnon-Bartsch, 2023), operating system, hardware and session details (Heil et al., 2021) and documentation(Ziemann et al., 2023). Software environments that use specific package versions can be controlled using containerization tools such as Docker (Merkel, 2014), or with package managers such as *renv* (Ushey & Wickham, 2025) for R, both integrated in the R *golem* (Fay et al., 2025) framework for Shiny applications.

While some interactive applications support the export of code, data, and reports(Barrett et al., 2013; Holscher et al., 2024; McMurdie & Holmes, 2015) in addition to language-specific data objects (Aussel et al., 2023) or environment containers (Torre et al., 2018), this information is typically organized in tool-specific ways. Our recently developed ENCORE framework is cross-disciplinary and guides researchers to structure and document their project in order to improve reproducibility and transparency and utilizes a standardized file system structure (sFSS) (van Kampen et al., 2024).

To enable an interactive exploration and analysis of T-cell and B-cell receptor repertoires measured using adaptive immune receptor repertoire sequencing (AIRR-Seq) (Boyd & Crowe, 2016; Freeman et al., 2009; Weinstein et al., 2009), we developed AbSolution (*manuscript under review*). AIRR-Seq analyses are particularly sensitive to both experimental parameters (e.g. sample preparation, library construction, and sequencing protocols) and analytical decisions (e.g. V(D)J annotation, error correction, and clonotype definition), as these directly affect clonal inference and diversity estimates, making rigorous documentation and reproducibility particularly critical (Breden et al., 2017; Peres et al., 2024). Although AbSolution offers an extensive interactive environment, it did not initially provide mechanisms to capture analysis provenance or ensure computational reproducibility. To address this, we integrated AbSolution with *shinymeta, golem*, and the ENCORE framework.

In this paper, we present our approach to enhance the reproducibility and transparency of interactive applications, using AbSolution as a proof-of-concept to address key issues in reproducible interactive analyses. The integrated application automatically generates a cross-disciplinary directory structure, a sFSS defined by the ENCORE framework, for organizing all files and documentation of a project, and provides automated export of all components required to reproduce an analysis, all organized according to the ENCORE structure. This self-contained export enables transparent sharing and reliable reproducibility of analyses. The reproducibility of this exported output has been independently validated by the CODECHECK initiative (Langton, 2025).

We conclude that systematic approaches addressing parameter and code capture and generation, software environments and containerization, and standardized file system structures are essential for achieving reproducibility in interactive applications. For Shiny applications, the combination of *shinymeta, golem* and ENCORE provides a robust and effective solution.

## Results

### AbSolution: an interactive tool for feature-based analysis and exploration of AIRR-seq data

AbSolution was developed to explore immune repertoires generated by AIRR-seq, with a particular focus on sequence-based features such as amino-acid biochemical properties (e.g., aliphatic index; (Ikai, 1980)) and sequence motifs. The application is implemented in R(R Core Team, 2021) and built using the Shiny package(Chang et al., 2024). An overview of the main analytical steps and the internal workflow is presented in *Figure 1*. The interface provides more than 50 configurable parameters, accessible through structured input controls (e.g., buttons, dropdown menus, sliders) or free-form inputs embedded within interactive visualizations.

**Figure 1.**
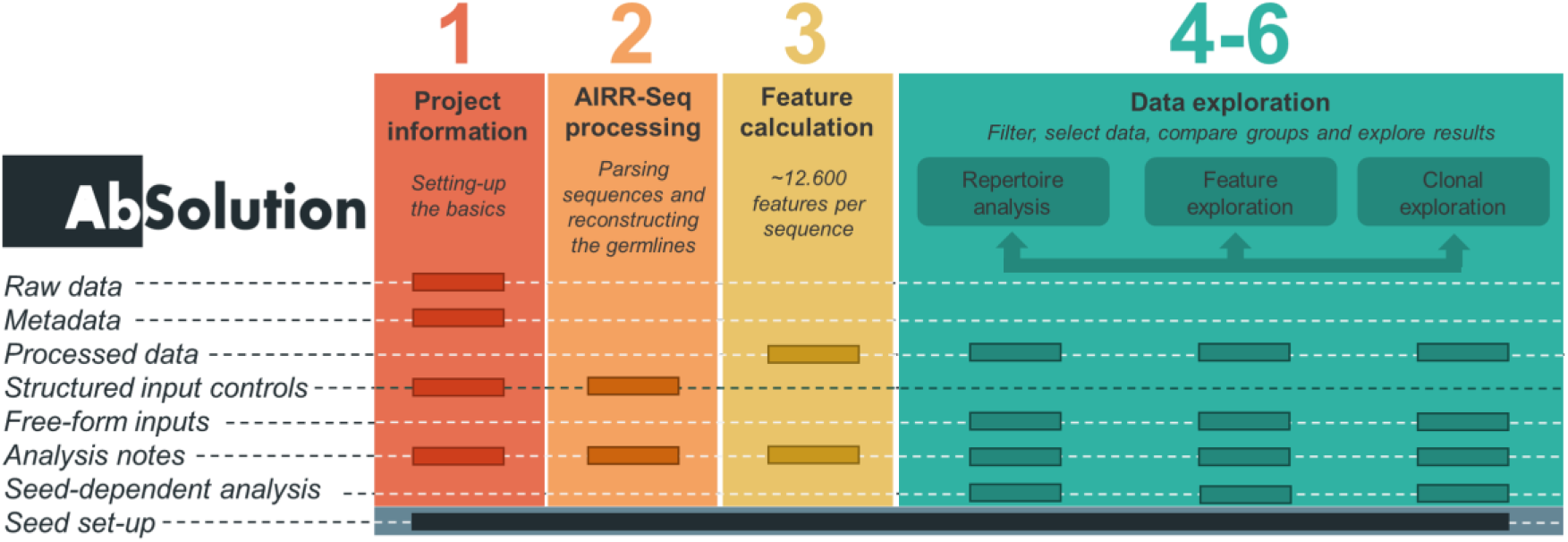
Schematic representation of the AbSolution workflow and input types. Steps 1-3 correspond to the preparatory and processing phases. (1) Project information: read (meta)data and general parameters (2) AIRR-Seq processing: sequence parsing and germline reconstruction. (3) Feature calculation: determination of 12,600 sequence-level features. Steps 4-6 include the analysis and exploration of repertoires (sequences), features and clones, in which processed data can be filtered, subsets selected, biological groups compared, and results examined. The lower section depicts the input and contextual elements that contribute to each step: raw data, metadata, processed data, structured input controls, free-form inputs, analysis notes, and presence of seed-controlled analyses. Figure adapted from AbSolution publication (manuscript under review).

### Data analysis and exploration

To demonstrate the reproducibility features of AbSolution, we analyzed a down-sampled dataset containing 1,393 V(D)J B-cell receptor sequences from CD19+ B cells isolated from the peripheral blood mononuclear cells of a healthy human donor, as provided in the *alakazam* R package (Gupta et al., 2015). After parsing and processing the dataset and its metadata in AbSolution (steps 1-3, *Figure 1*), the application enables the analysis and exploration of sequence-based repertoire features across the complementarity-determining regions (CDR1-3) and framework regions (FRW1-3) of the sequences(Lefranc et al., 2003) and of their unmutated common ancestor sequences (germline sequence) (*Figure 2*).

**Figure 2.**
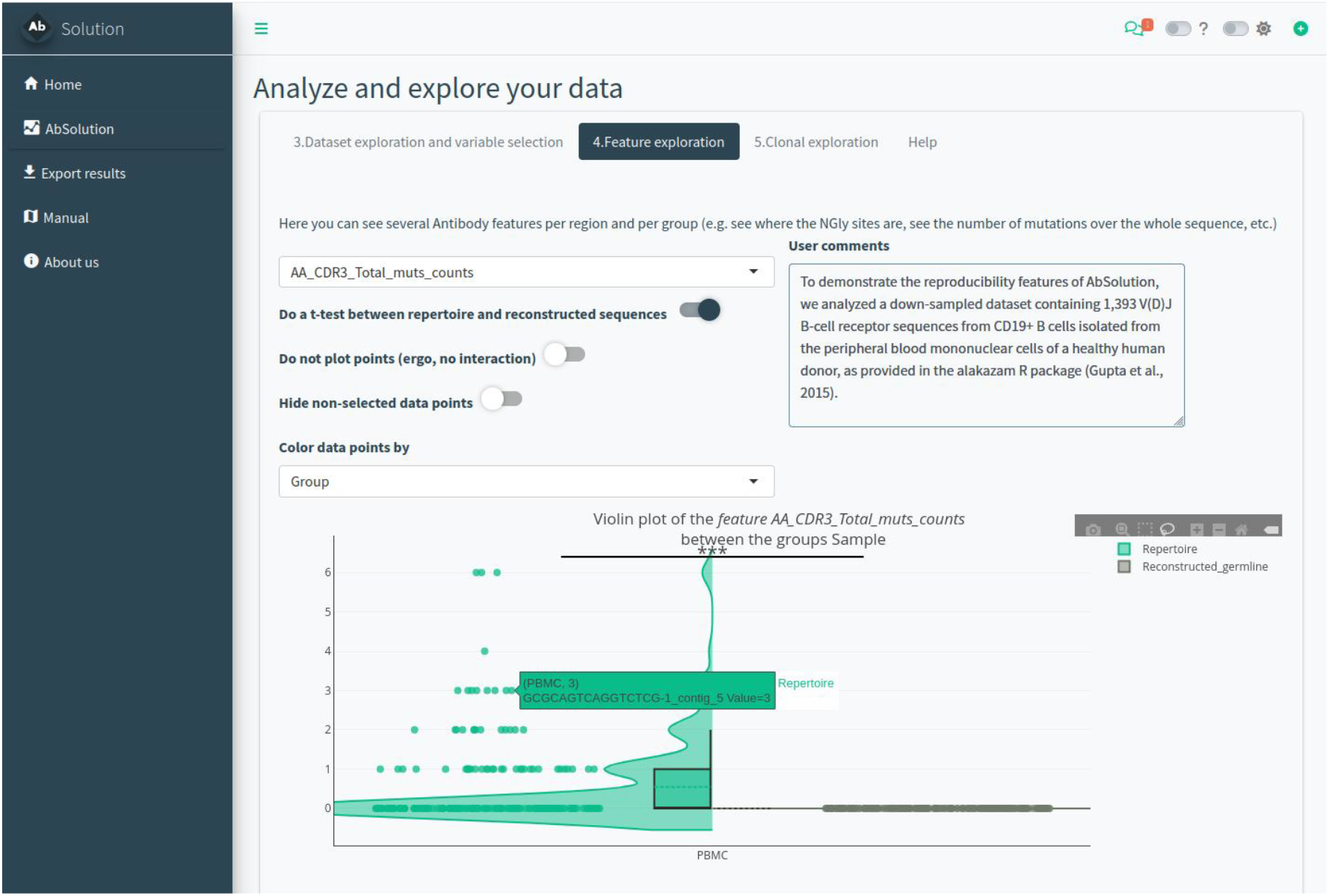
Screenshot of AbSolution. This page shows an interactive split-violin plot of the number of mutations in the CDR3 region of the repertoire sequences and reconstructed germline sequences. Parameters to generate this plot and a comment for this specific exploration step are shown above the plot.

All sequence, feature and clonal visualizations and summaries update automatically in response to the choices made during the analyses. These choices include filtering criteria (e.g., removal of highly and/or lowly mutated sequences), selection of specific subsets (e.g., sequences with a specific V(D)J gene usage), and definition of biological groups for comparison (e.g., donor subsets or experimental conditions). Parameter choices in the various analyses steps directly determine the results, and being able to store their values is critical for ensuring both reproducibility and accurate interpretation.

### Implementation of a framework to reproduce interactive analysis and exploration

To support reproducibility of AbSolution we integrated three complementary framework elements: i) the capture of domain logic from interactive operations, ii) a controlled and portable software environment, and iii) a standardized structure for both the organization of the project and the export of complete analytical outputs.

#### i) Capturing domain logic within interactive analyses

Interactive analysis make it hard to exactly reproduce results unless the underlying analytical decisions and parameters are explicitly captured. AbSolution addresses this challenge by capturing the domain logic in reactive operations and converting it into executable R code. This has been enabled through the integration of the *shinymeta* package, which encapsulates internal reactive expressions and variables into explicit code-generating segments. These code segments are then interlinked within a unified R markdown notebook, which can be executed independently of AbSolution to reconstruct the full analysis and results. This guarantees that the interactive workflow is rendered as a transparent, ordered, and reproducible computational sequence; however, it also requires that reproducibility support be explicitly implemented modifying the source code, and the required effort and complexity increases with the size and functional scope of the application.

#### ii) Ensuring a fully reproducible software environment

Reproducing an interactive data analysis requires a fixed software environment containing software and package versions used during the analyses. AbSolution was implemented as a *golem*-based application to enforce modularity and systematic version control of the application and its dependencies through *renv* and Docker. A local snapshot of all package versions is made using *renv* to ensure that analyses can be executed with the exact same package versions outside containers. The entire environment, operating system, R and package versions is provided as a Docker image to allow the full analysis pipeline to be executed identically on different operating systems and hardware platforms and at future points in time.

#### iii) Standardized structure for project organization

AbSolution adopts the ENCORE framework, which is based on a standardized file structure (compendium) and documentation templates to organize and describe (meta)data, code, results, and project information (van Kampen et al., 2024). This not only contributes to reproducibility but also transparency of a data analysis project. Moreover, the ENCORE structure facilities easy sharing of complete projects with peers or archiving projects in repositories such as Zenodo. To support the implementation of an ENCORE-compliant environment for any given project, AbSolution enables direct download and setup of the latest ENCORE template (*Figure 3*) from GitHub, which acts as a standardized file system structure in which raw and meta data are stored in the *Data* folder, while the *Processing* directory serves as the designated workspace for running analyses. All exported analysis is typically stored in the *Processing* directory of the project compendium.

**Figure 3.**
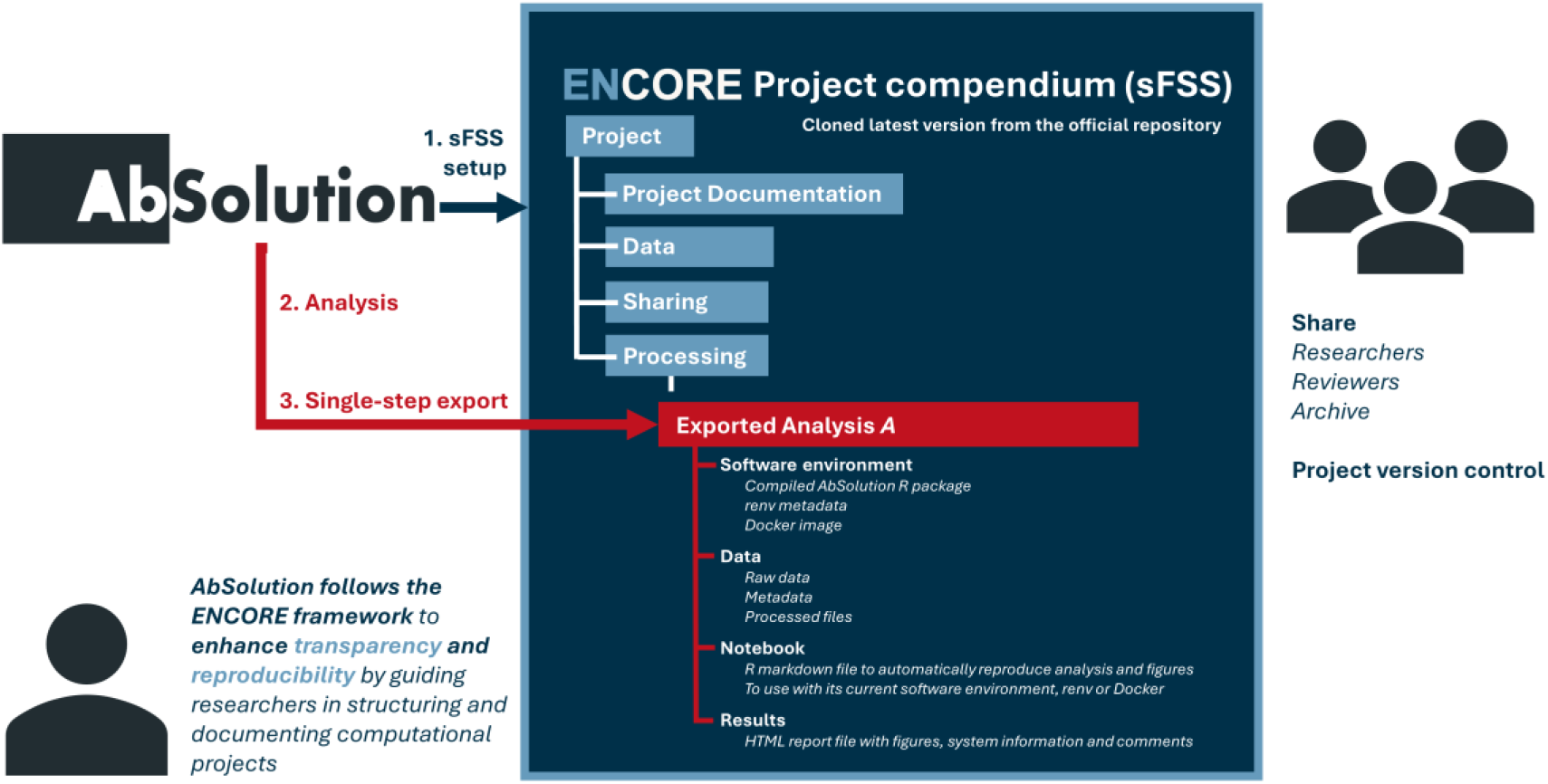
Overview of AbSolution following the ENCORE framework. The standardized file system structure (sFSS) serves as the scaffold for new projects, and is created by cloning the template from the ENCORE GitHub repository when starting a project. AbSolution automates the generation of the base sFSS for the researchers. Raw and meta data have to be stored in the Data folder, while the Processing directory serves as the designated workspace for running analyses. After completing data analysis and exploration with AbSolution, a self-contained ZIP archive can be exported and stored in the Processing directory of the ENCORE project compendium, containing all files required to reproduce the analysis. Note that ‘Analysis A’ is used as an example and a more descriptive name is given in practice. The archive includes the software environment (AbSolution package version, Dockerfile, and renv metadata), the full dataset (raw AIRR-C files, metadata, and processed intermediate files), an R Markdown notebook that includes inputs, parameters and notes and reproduces the workflow using relative paths, and an HTML report with inputs, parameters, notes, generated figures, operating system, hardware and R session details, and package versions, thereby enabling transparent, shareable, and fully reproducible computational analyses. Figure adapted from the ENCORE publication (van Kampen et al., 2024).

### Automating the export of a standardized, structured, and self-contained reproducible output

After finishing data exploration with AbSolution, a self-contained zip archive can be exported can be exported containing all relevant files organized according to the ENCORE structure and intended for storage in the *Processing* directory of the project compendium. The zip archive contains the following subfolders, with some being optional to minimize redundancy and allow for flexibility:

- **/0_SoftwareEnvironment/R**. Contains the AbSolution package archive (tar.gz) corresponding to the exact version used during the analysis, supporting local installation independent of external repositories, a Docker file defining the complete containerized computational environment, and the *renv* metadata (Ushey & Wickham, 2025), thereby providing alternative, complementary strategies for reproducing the analysis depending on the preferred computational setup.
- **/Data/Dataset**. Contains the dataset used for the analysis.
  - **/Raw**. AIRR-Seq repertoire sequence files in AIRR-C format (Vander Heiden et al., 2018) for each sample.
  - **/Meta**. Metadata file containing sample information.
  - **/Processed**. Intermediate files with sequence and feature information produced during the analysis. These files are used for the generation of the main results and required for data exploration.
- **/Notebook**. Contains a R markdown file used to reproduce the analysis and generate the figures. Use of relative paths ensure the execution of the notebook. For efficiency, markdown chunks responsible for generating the processed files are not evaluated when generating the zip archive. Analysis and parameter details, contact information, and other notes incorporated during the interactive analysis are included in the notebook. The R markdown file serves as a precise record of the analytical choices made during the session and can be used as a reference to repeat the same analysis within the application. In addition, the exported code enhances transparency and facilitates reproducibility, allowing complete analyses to be shared, inspected, re-executed and followed-up by other researchers. The alternative environment specifications in /0_SoftwareEnvironment/R enable users to reproduce the R Markdown notebook using the setup best suited to their local constraints. The resulting outputs can then be compared against the reference HTML report in /Results to validate the reproduced analysis.
- **/Results**: Includes the HTML report file (*Figure 4*) generated from the R markdown file, containing the figures generated with AbSolution. These figures can be saved following the graphical configuration (format, dimensions and scale) set by the author of the analysis. The HTML file includes comprehensive operating system, hardware, R session and package versioning details.

**Figure 4.**
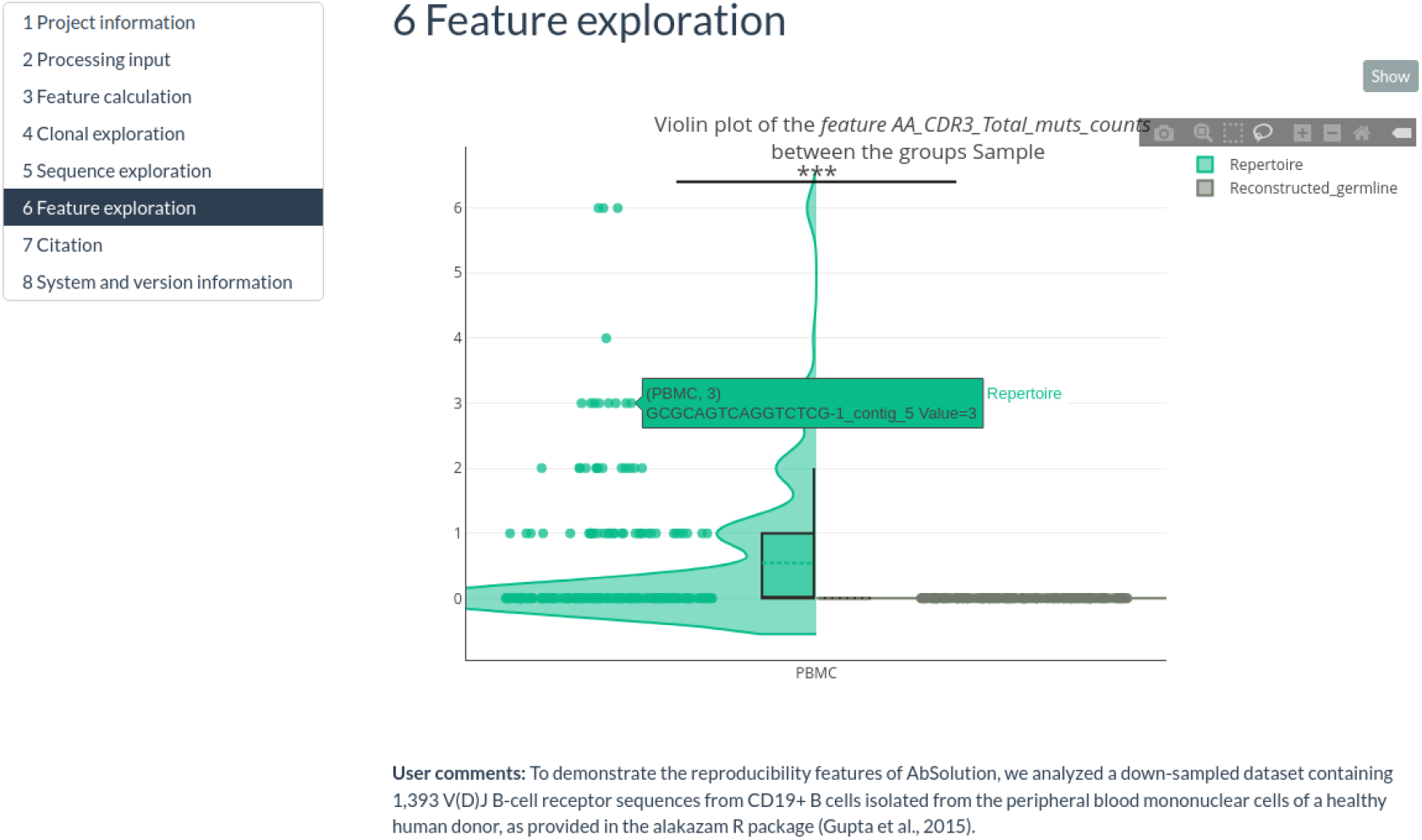
Exported HTML report from the markdown Notebook. This interactive split-violin plot of the number of mutations in the CDR3 is identical to the graph in AbSolution (Figure 2) but has been generated using domain-logic capture in the application. This code is hidden in the report, but can be shown from the Code button. A comment that was saved during analysis is shown below the plot.

### Independent validation of reproducibility

To externally verify reproducibility, the analysis output produced by AbSolution was subjected to a formal CODECHECK review. CODECHECK (Nust & Eglen, 2021) is an external reproducibility service in which a verifier re-runs the computational workflow using the provided exported files and documentation, checking that all steps execute as described and that the reported results can be independently produced. For this study, the verifier successfully reproduced all figures and outputs from the HTML report file with “no errors first time”, as documented in the CODECHECK certificate associated with this paper (Langton, 2025) (https://zenodo.org/records/16758977).

## Discussion

Ensuring transparency and reproducibility of computational analyses remains a persistent challenge across scientific domains, and these challenges are amplified in the context of interactive applications. Unlike scripted workflows, interactive analyses involve manual exploratory steps that are typically not captured, making it difficult to recover the full analytical provenance. Achieving reproducibility therefore requires not only access to data and code, but also recording and storage of all inputs, executed logic, software environments, containerized environments, researcher annotations, and operating system, hardware and session details.

In this work, we demonstrate that computational reproducibility of a complex interactive application can be achieved by integrating domain capture logic (*shinymeta* (Cheng & Sievert, 2025)), management of software environment (*golem* (Fay et al., 2025)), and the ENCORE framework(van Kampen et al., 2024). The resulting export contains all components needed to reproduce the analysis -even across systems-. The reproducibility of our proof-of-concept has been confirmed through independent validation through a CODECHECK assessment (Langton, 2025). Importantly, our approach automates the entire compilation and export procedure and enables reproducibility beyond code availability by also making the interactive analysis and exploration steps explicit in a R markdown notebook; a dimension often overlooked in existing reproducibility frameworks.

Nevertheless, our proof-of-concept highlights several current limitations and areas for improvement. First, although *shinymeta* greatly facilitates the capture of reactive logic, adapting complex pipelines still requires nontrivial developer effort and training, particularly when the workflow spans multiple modules and reactive expressions. Second, while ENCORE provides a general-purpose structure, domain-specific adjustments may be needed to accommodate software or metadata standards, for instance, adjusting the layout to conform to the consensus structure of an R package. Third, long-term reproducibility remains sensitive to evolving software ecosystems; even with containers, updates in underlying infrastructures or discontinuation of packages may eventually require maintenance. Finally, although the export process is automated, researchers must still ensure appropriate annotations and responsible tool and data handling for reproducibility to be fully effective. Despite these limitations, we demonstrated that a standardized, automated framework can markedly reduce the obstacles to reproducibility in interactive applications. It offers a concrete design pattern and reference implementation for capturing reactive logic as executable notebooks, managing dependencies, and structuring outputs in a principled way, moving beyond existing tools that export only partial scripts or static reports and typically leave the organization of reproducible artefacts to project-specific solutions. Future work should focus on extending this framework to interactive applications beyond Shiny (e.g., Streamlit in Python), harmonizing output format with research data repositories such as Zenodo to advance FAIR compliance, and fostering community consensus toward a standardized, application-agnostic reproducibility model.

Overall, by providing an extensible and field-agnostic solution, our proof-of-concept represents a step forward toward reproducible science when using interactive applications.

## Materials and methods

### Software

AbSolution is an R package publicly accessible in CRAN (https://cran.r-project.org/package=AbSolution) for installation. It provides an R Shiny interface (Chang et al., 2024) to facilitate exploratory data analysis. The package supports the analysis of large-than-RAM datasets through by using memory-mapped matrices implemented through the *bigstatsr* package (Prive et al., 2018) and by leveraging linear-scaling algorithms whenever possible. AbSolution adheres to the ENCORE workflow for reproducible computational research (van Kampen et al., 2024). The complete ENCORE development folder for this project is available at https://github.com/EDS-Bioinformatics-Laboratory/AbSolution. The package version used for the CODECHECK report can be found under the commit id ca26697.

### Hardware

For the data analysis, AbSolution was executed in Ubuntu 18.04.6 LTS, Intel® Core™ i7-8750H CPU @ 2.20GHz × 12 GB RAM. For the CODECHECK reproduction, the analysis was executed in Windows 11, Intel® Core™ i5-8250U CPU @ 1.60GHz (1.80 GHz).

## Acknowledgements

This work is supported by COSMIC which has received funding from the European Union’s Horizon 2020 research and innovation programme under the Marie Skłodowska-Curie grant agreement No 765158. Authors thank Huub Hoefsloot for his support in the development.

## Contributions

R.G.V. and Av.K developed AbSolution and ENCORE and integrated both approaches. S.L. independently executed the exported code and confirmed its reproducibility through the CODECHECK workflow. All authors contributed to the writing of the manuscript.

## Data availability

The dataset analysed during the current study is publicly available from the *alakazam* (Gupta et al., 2015) R package, under the *Example10x* command. The exported reproducible output of the AbSolution analysis presented in this paper is available from Github (https://github.com/codecheckers/absolution_codecheck).

## Code availability

Source code used for this work is available from GitHub [https://github.com/EDS-Bioinformatics-Laboratory/AbSolution]. sFSS used for AbSolution is available from Zenodo [https://github.com/EDS-Bioinformatics-Laboratory/ENCORE]. CODECHECK certificate confirming that the computations underlying this article could be independently executed is available from Zenodo [https://zenodo.org/records/16758977].

